# Immune-challenged vampire bats produce fewer contact calls

**DOI:** 10.1101/2020.04.17.046730

**Authors:** Sebastian Stockmaier, Daniel. I. Bolnick, Rachel. A. Page, Darija Josic, Gerald G. Carter

**Author notes:** Corresponding author: Sebastian Stockmaier –.

## Abstract

Infections can affect how animals vocalize and, hence, interact with conspecifics. While this effect has been shown for mate-attraction calls, other vocalizations that facilitate social contact have received less attention. When isolated, vampire bats produce contact calls which attract highly associated groupmates. Here, we test the effect of an immune challenge on contact calling rates of individually isolated vampire bats. Sickness behavior did not appear to change call structure, but it decreased the number of contact calls produced. This effect could decrease contact with groupmates and augment other established mechanisms by which sickness reduces social encounters (e.g. mortality, lethargy, and social withdrawal or disinterest).

## Introduction

Infections can reduce contact between individuals by inducing ‘sickness behavior’. For instance, sickness can decrease physical social encounters through reduced movement (Lopes et al., 2016) or decrease directed social interactions like grooming (Stockmaier et al., 2020, 2018). Reductions in social contact can also occur if infected individuals vocalize less. For example, an immune challenge reduces male mate-attraction vocalizations or some of their components in several species (Dreiss et al., 2008; Fedorka and Mousseau, 2006; Garamszegi et al., 2004; Jacot et al., 2004; Lopes and König, 2016; York et al., 2016). If sick males attract fewer females and if avoiding sick males decreases the likelihood of females acquiring parasites (Hillgarth, 1996; Luong et al., 2000; Martinez-Padilla et al., 2012) then transmission of parasites between the sexes will decrease (Able, 1996).

Besides courtship vocalizations, a broader range of vocal interactions could be influenced by sickness, but these other call types have received less attention. In many group-living animals that live in conditions of low visibility, or that must maintain cohesion while on the move, individuals produce contact calls to maintain contact with groupmates or particular affiliated individuals (Arnold and Wilkinson, 2011; Bradbury and Vehrencamp, 1998; Carter et al., 2012, 2009; Carter and Wilkinson, 2016; Chaverri et al., 2010; Cortopassi and Bradbury, 2006; Janik and Sayigh, 2013; Maurello et al., 2000; van Oosterom et al., 2016). If contact calls facilitate physical contact, then sickness behavior that reduces the rate of contact calling should decrease contact with groupmates. However, if contact calling is used by an individual in need to gain benefits from others, then sick individuals might instead make a greater number of contact calls. For example, when parents can acquire enough food to feed all their offspring, hungry nestlings in worse condition are expected to call more often, not less (Caro et al., 2016; Godfray, 1991). The expected effect of sickness behavior on contact calling by distressed individuals is therefore less clear.

Isolated common vampire bats (*Desmodus rotundus*) produce multi-harmonic contact calls that vary in spectral structure and duration, and that facilitate individual contact and recognition (Carter et al., 2012; Carter and Wilkinson, 2016, Fig. 1B, C). Contact calls appear to be important for maintaining co-roosting associations with bonded partners and for finding or recruiting those partners for help; for example, trapped and hungry individuals appear to use contact calls to recruit both kin and non-kin food donors to feed them by regurgitation (Carter et al., 2017; Carter and Wilkinson, 2016).

**Figure 1.**
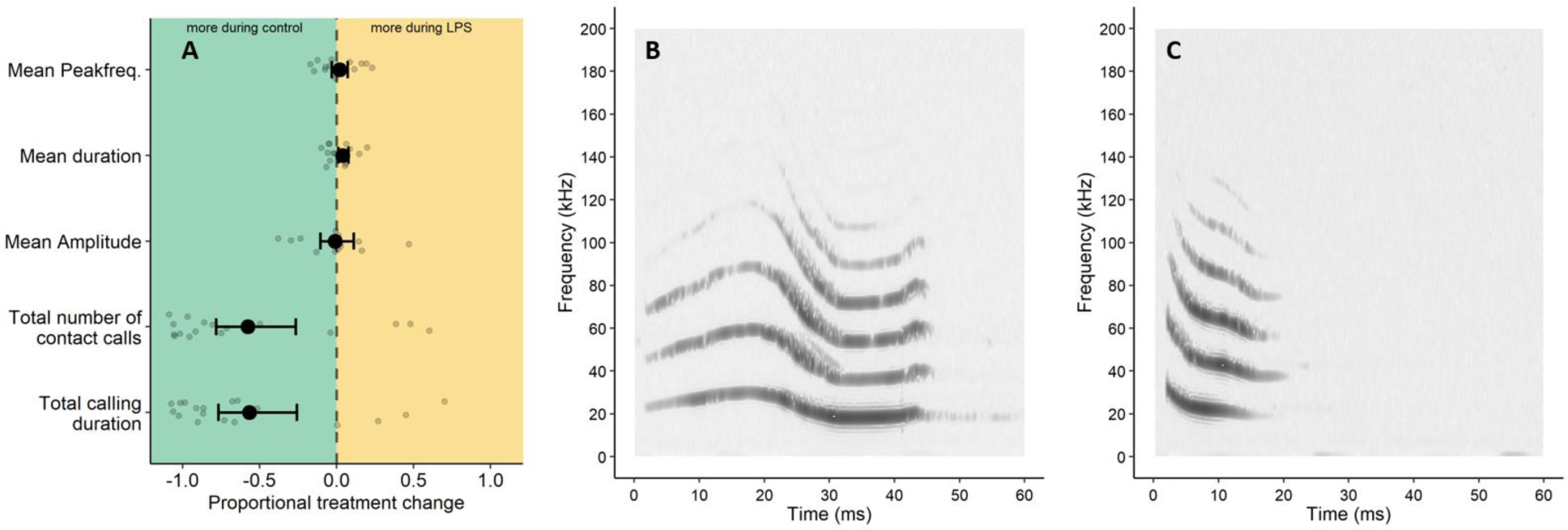
Effect of LPS on vampire bat contact calls. **Panel (A)** shows mean standardized LPS-effect ± bootstrapped 95% confidence intervals (see Table S1) and data points for the mean peak frequency, mean duration, mean amplitude, the total number of contact calls produced, and the total calling duration. Example spectrogram of a longer **(B)** and shorter **(C)** vampire bat contact call that also vary in peak frequency (darker regions show frequencies of greater relative amplitude).

Here, we mimicked a bacterial infection in vampire bats by using lipopolysaccharide (LPS) to trigger transient physiological symptoms and sickness behaviors (Ripperger et al., 2020; Stockmaier et al., 2020, 2018), and then we test for the effect of LPS on contact calling behavior. LPS-injected vampire bats are groomed by fewer bats and have lower social connectedness in the wild, an effect that could be driven in part by a reduction in contact calling (Ripperger et al., 2020; Stockmaier et al., 2020). We show that LPS-induced sickness behavior decreases the number of contact calls produced by isolated vampire bats. This effect is relevant for pathogen transmission in social animals that rely on vocalizations to maintain contact because it might further reduce the probability of physical contact between individuals beyond the effects of reduced movement.

## Material and Methods

We recorded contact calls by physically isolating an adult female vampire bat (n = 18) in a soft mesh cage at a distance of 10 - 30 cm from a CM16 ultrasound condenser microphone (frequency range 1-200 KHz, Avisoft Bioacoustics, Berlin, Germany). The mesh cage was inside a 68-liter plastic bin lined with acoustic dampening foam and within hearing range of conspecifics of a captive colony. To selectively record contact calls, we used a digitizer (116 Hn UltrasoundGate, Avisoft Bioacoustics, Berlin, Germany, sampling frequency of 250 or 500 kHz) to save a .wav file whenever a 10-50 kHz sound was detected at greater than 5% amplitude. We used Avisoft SASLabPro (Avisoft Bioacoustics, Berlin, Germany) to measure the onset, duration, peak amplitude, and peak frequency of all calls. We excluded echolocation calls and other noise by deleting sounds that were longer than 60 ms and shorter than 10 ms.

For each bat, we recorded two trial types. In LPS trials, we induced sickness behavior in subjects by injecting them subcutaneously with lipopolysaccharide (LPS, L2630 Sigma-Aldrich, USA, dose: 5 mg/kg body mass of bat) in phosphate-buffered saline (PBS) before the recording period. We chose this dose based initially on observed effects in another bat species (Stockmaier et al., 2015), and on later studies in vampire bats, which showed that this dose increases white blood cell count and neutrophil to lymphocyte ratio (Stockmaier et al., 2018) and decreases in physical activity, social encounters, and social grooming (Ripperger et al., 2020; Stockmaier et al., 2020, 2018). In control trials, the same bats were injected with an equivalent volume of only PBS as a control treatment.

Treatments were given in random order, and 8 bats received the control treatment first. We recorded bats for 3-5 hours immediately after the injection, because we previously detected symptoms for at least 6 hours post-injection (Stockmaier et al., 2018). Different bats were recorded for different times after injection, but the paired LPS and control trials were always the same duration and time of the night. The inter-trial period was at least 5 days to ensure recovery of the bats (Ripperger et al., 2020; Stockmaier et al., 2018). To calculate a standardized effect (proportional change) of LPS on vocalizations for each bat, we used (Y_LPS_ - Y_C_)/(Y_LPS_ + Y_C_), where Y_LPS_ and Y_C_ are the measures of vocal activity during the bat’s LPS and control trial, respectively.

To test for an effect of LPS on contact calling, we randomly swapped the control and LPS trial data within each bat to calculate a distribution of t-statistics under the null hypothesis of no difference between the LPS and control trial, then compared the observed t-statistic to this distribution to obtain a two-sided p-value (i.e. a nonparametric permuted paired t-test). To estimate 95% confidence intervals for LPS effect sizes, we used nonparametric bootstrapping with accelerated bias-corrected percentile limits (Puth et al., 2015). We used 5000 permutations for both methods. We calculated the mean and bootstrapped 95% CI for the LPS effect on five measures of contact calling behavior: the total number of contact calls produced, the sum of call durations, the mean call duration, the mean amplitude, and the mean peak frequency (the frequency at the point of the maximum amplitude of the entire element). Our data and the R script are available on Figshare (https://doi.org/10.6084/m9.figshare.11861877.v3). Our work was approved by the Smithsonian Tropical Research Institute Animal Care and Use Committee (#2016-0728-2019-A2) and the Animal Care and Use Committee of the University of Texas at Austin (AUP-2016-00124).

## Results

LPS injections led to fewer contact calls. The average contact calling rates per bat during the control and LPS trials were respectively 66 and 16 contact calls per hour (see Fig. S1 for details on each bat). On average, LPS injections caused female vampire bats to produce 30% fewer contact calls, with 15 of 18 bats producing fewer contact calls during the LPS trial compared to the control trial (p = 0.0006, Fig. 1A, Table 1, Fig. S1). Fewer calls led to an average decrease of 32% in total calling duration (p = 0.0012, Fig. 1A, Table 1). We did not detect an effect of LPS on mean call amplitude (Fig. 1A, Table 1). Although vampire bats produce contact calls that vary in call structure (Fig. 1B, C), we did not detect an effect of LPS injections on mean call duration (Fig. 1A, Table 1) or mean peak frequency (Fig. 1A, Table 1).

**Table 1:** Effect of LPS on vampire bat contact calls. Means and their bootstrapped 95% confidence intervals for the standardized LPS effects for contact call number, total calling duration, mean amplitude, mean peak frequency, and mean call duration (Table corresponding to **Figure 1A**).

| Measure | N | Lower CI | Mean | Upper CI | p-value |
| --- | --- | --- | --- | --- | --- |
| Call number | 18 | -0.78 | -0.58 | -0.27 | 0.0006 |
| Calling duration | 18 | -0.77 | -0.57 | -0.26 | 0.0012 |
| Mean amplitude | 14* | -0.11 | -0.01 | 0.11 | 0.8754 |
| Mean peak frequency | 14* | -0.03 | 0.02 | 0.07 | 0.4656 |
| Mean duration | 14* | 0.01 | 0.04 | 0.08 | 0.0510 |

\* We could not calculate the LPS effect on mean amplitude, mean peak frequency and mean duration for four bats because they did not produce any calls when injected with LPS.

## Discussion

Infection-induced sickness behaviors can affect vocal communication as evident in LPS-injected male house mice that produce fewer call syllables which reduces associations with females (Lopes and König, 2016), and immune-challenged males decreasing their song rate in collared flycatchers, white-browed sparrow weavers, field crickets, and white-crowned sparrows (Garamszegi et al., 2004; Jacot et al., 2004; Owen-Ashley et al., 2006; York et al., 2016). In comparison to these mate-attraction calls, contact calls and signals of need are interesting to consider because a state of poor condition could lead to either a higher or lower calling rate. In vampire bats, contact calling can attract food donors and might act as a signal of need (Carter et al., 2012; Carter and Wilkinson, 2016). Here, we showed that an immune challenge reduces contact calling, which could potentially help to explain why immune-challenged vampire bats encounter fewer individuals (Ripperger et al., 2020; Stockmaier et al., 2020), in addition to the more obvious explanation of reduced movement.

LPS-injected vampire bats spend less time awake, moving, or grooming (Stockmaier et al., 2018), so our results are most consistent with the simplest explanation that reduced contact calling is also due to lethargy as opposed to a kin-selected behavior of active self-isolation, which has only been observed in eusocial insects (Bos et al., 2012; Heinze and Walter, 2010; Stroeymeyt et al., 2018). We hypothesize that vampire bats reduce contact calling to support the energetic demands of the physiological response. Across several taxa, the metabolic costs of acoustic signaling are estimated to be about eight times that of remaining silent (Ophir et al., 2010) and call rate is sensitive to other ecological constraints like reduced food availability (Ritschard and Brumm, 2012). Experiments with a related frugivorous bat species show that LPS injections reduce body mass and increase resting metabolic rate by 40% (Guerrero et al., 2018).

We used a dose of LPS for which we knew the physiological and behavioral effects in vampire bats (Ripperger et al., 2020; Stockmaier et al., 2020, 2018). It is important to note, however, that the physiological responses to LPS are dose dependent and involve both pro-inflammatory and anti-inflammatory responses (Armour et al., 2020). To determine what doses are most ecologically relevant for different diseases, future work must compare the relationship between the dose-dependent effects of LPS against the effects of natural bacterial infections in vampire bats and other species.

It is also important to note that infection-induced changes to social vocalizations are pathogen-specific. LPS mimics common symptoms of a bacterial infection in vampire bats and other animals (Lopes et al., 2016; Stockmaier et al., 2018). Some live pathogens, however, could increase specific social behaviors to favor their transmission (Klein, 2003). For instance, chytrid-fungus infected Japanese tree frogs (*Hyla japonica*) increase their mating call effort which potentially favors the transmission of the fungus (An and Waldman, 2016). Vampire bats harbor multiple pathogens in their saliva that rely on directed social interactions, like Bartonella (Becker et al., 2018), hemoplasmas (Volokhov et al., 2017), and most notably, rabies (Aguilar-Setien et al., 2005). It would be particularly interesting to look at how rabid vampire bats change their calling behavior and their response to calls of conspecifics.

Besides call rate, the structure of animal vocalizations might also depend on infection status (Dreiss et al., 2008; Fedorka and Mousseau, 2006). Common vampire bats produce highly variable contact calls (e.g. Figure 1 B and C), but we found no evidence that sick bats consistently produced any particular contact call structure more or less. However, for some pathogens such as rabies, which could affect the vocal tract there may be clear differences in call structure.

## Acknowledgements

We thank the Smithsonian Tropical Research Institute for logistical support. Samuel Kaiser, Vanessa Pèrez, Jineth Berrio-Martinez, Imran Razik, Bridget Brown, David Girbino, Simon Ripperger, and Emma Kline for help in the field and animal caretaking.

## Author contributions

SS and GGC designed the study and carried out the experiments and husbandry. GGC captured the bats and established the captive colony. SS, GGC and DJ contributed to the data collection and analysis. GGC, DIB, and RAP coordinated the study and provided valuable resources and lab space. All authors contributed to draft the manuscript and gave final approval for publication.

