## Supplementary material for "Immune-challenged vampire bats produce fewer contact calls": Fig. S1

### Figure S1

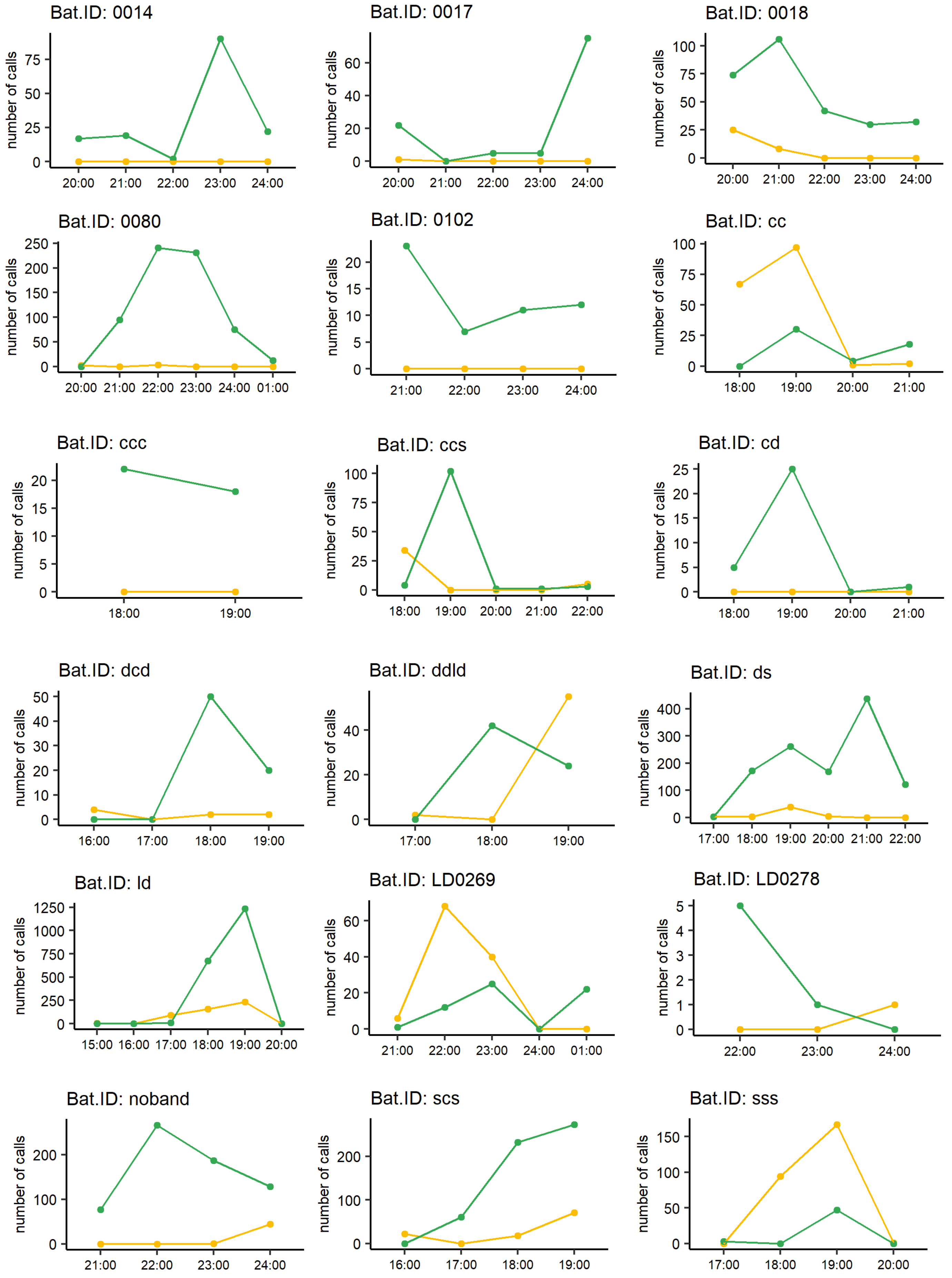


**Figure S1:** Plots show comparisons of total number of calls produced per 60-min time bin for the same bats after LPS (yellow) and control (PBS, green) injections. A datapoint at 21:00 counts calls from 21:00 to 22:00
